# A conserved Foxp1-Pde4d axis regulates inflammatory responsiveness

**DOI:** 10.64898/2026.10.27.762160

**Authors:** Loes M.H. Elemans, Panna Tandon, Gavin Rodger, Lauren Stirling, Samantha Hendry, Muhammad Nadeem Akhtar, Xinyi Chen, Tijana Mitić, Yi Feng, Robert K. Semple, James E.N. Minchin

**Affiliations:** Institute for Neuroscience and Cardiovascular Research, University of Edinburgh, Scotland, UK; Institute for Regeneration and Repair, University of Edinburgh, Scotland, UK; Cancer Research UK Scotland Centre, Institute of Genetics and Cancer, University of Edinburgh, Scotland, UK; Institute of Genetics and Cancer, University of Edinburgh, Scotland, UK

## Abstract

Inflammatory responses are heterogeneous, even among cells exposed to a common insult, yet the regulatory mechanisms that distinguish these inflammatory states remain poorly understood. Here, we leveraged inflammatory state heterogeneity to identify a conserved Foxp1-Pde4d regulatory relationship and uncover a role for this axis in preserving inflammatory responsiveness. Genotoxic stress in adipocyte progenitors generated heterogeneous engagement of type-I interferon (IFN) and NFκB transcriptional programmes, and transcriptome-wide analysis independently identified Foxp1 and the cAMP phosphodiesterase Pde4d among genes associated with lower IFN- and higher NFκB-associated transcription. ChIP-seq revealed selective loss of Foxp1 occupancy at a Pde4d-associated regulatory element following genotoxic stress, while Foxp1 gain- and loss-of-function established Foxp1 as a positive regulator of Pde4d expression. Remarkably, coupling of Foxp1-Pde4d expression was broadly conserved across vertebrate tissues and damage models. In vivo, Foxp1 loss in zebrafish revealed a striking dissociation between inflammatory state and response capacity: basal NFκB activity was elevated, yet injury-induced NFκB activation was markedly blunted and macrophage recruitment impaired. Adenylyl cyclase inhibition restored macrophage recruitment in Foxp1-deficient zebrafish, functionally linking the Foxp1 phenotype to cAMP signalling and supporting a model in which Foxp1-Pde4d regulation preserves inflammatory responsiveness through control of cAMP. Together, these findings show how inflammatory-state heterogeneity can reveal conserved regulatory mechanisms linking basal cellular state to subsequent response capacity.

## Introduction

Effective inflammatory responses require continuous adaptation to changing cellular and tissue conditions (Natoli and Ostuni 2019). Inflammatory pathways must be sufficiently active to coordinate immune surveillance and tissue repair, yet appropriately constrained to avoid persistent inflammation and tissue damage (Wong and Germain 2018). Inflammatory responses are also heterogeneous, with cellular state, signalling history and the temporal sequence of stimuli shaping subsequent responsiveness (Lei and Tsang 2025). How such heterogeneity is regulated, and whether the mechanisms that establish inflammatory state also determine the capacity to respond to subsequent challenge, remain poorly understood.

DNA damage provides an important context in which to investigate this problem. Genotoxic stress accumulates throughout life and contributes to ageing, cancer and chronic inflammatory disease (Siametis and Garinis 2025). Persistent DNA damage can engage innate nucleic-acid sensing pathways, including cGAS-STING, triggering inflammatory and antiviral transcriptional programmes (Dou et al. 2017; Glück et al. 2017; Decout et al. 2021; Gulen et al. 2023). Prominent among these are type-I interferon (IFN) and NFκB, which can arise downstream of common damage-sensing pathways but are mechanistically distinct and need not be engaged synchronously (Yu et al. 2015; Rodier et al. 2009). IFN establishes an antiviral-like transcriptional state and can reinforce cell-cycle arrest, whereas NFκB drives inflammatory secretory programmes involved in immune-cell recruitment and tissue remodelling (Frisch and MacFawn 2020; Wang et al. 2024). Thus, genotoxic stress provides a setting in which a common cellular insult can generate distinct inflammatory programmes, but the intrinsic mechanisms that determine their relative engagement remain poorly understood.

Cyclic AMP (cAMP) is well positioned to contribute to such inflammatory-state regulation. Generated by adenylyl cyclases and degraded by phosphodiesterases (PDEs), cAMP can promote or restrain inflammatory signalling depending on cellular and signalling context (Gerlo et al. 2011; Sheng et al. 2025; Qiang et al. 2018). PDEs are central to this context dependence, controlling the spatial and temporal organisation of cAMP signals and thereby shaping local signalling thresholds (Baillie 2009; Houslay 2010). Different cell types rely on distinct PDE family members: PDE4B is an established regulator of inflammatory signalling in immune cells (Jin and Conti 2002), whereas PDE4D is prominent in mesenchymal and adipose tissues, where it controls compartmentalised cAMP signalling (Krier et al. 2025; Seigner et al. 2025; Huang et al. 2021; Meng et al. 2017). Regulation of PDE expression could therefore provide a mechanism through which cellular state shapes both basal signalling and the capacity to respond to subsequent challenge. Whether this regulatory layer is remodelled by cellular stress, and whether such remodelling contributes to inflammatory responsiveness, remain unknown. Known transcriptional regulators of PDE expression predominantly participate in feedback circuits downstream of cAMP signalling itself (Ke et al. 2015; Zhang et al. 2019; Soda et al. 2013), leaving it unclear how broader changes in cellular state are communicated to the cAMP regulatory machinery. Identifying transcriptional mechanisms that link cellular stress to PDE expression could therefore reveal how inflammatory state and responsiveness are recalibrated as tissue conditions change.

Here, we use genotoxic stress in adipocyte progenitors to investigate the regulatory basis of inflammatory-state heterogeneity. Single-cell profiling identified Foxp1 and Pde4d as strongly associated with the relative engagement of IFN- and NFκB-associated transcriptional programmes. We show that Foxp1 positively regulates Pde4d expression through a damage-sensitive transcriptional relationship that is deeply conserved across vertebrates. Extending these findings in vivo, Foxp1 loss elevated basal NFκB activity while impairing its subsequent induction and macrophage recruitment following tissue injury, with recruitment restored by adenylyl cyclase inhibition. Together, these findings identify a conserved Foxp1-Pde4d regulatory axis associated with inflammatory-state heterogeneity and reveal a role for this axis in preserving inducible inflammatory responsiveness.

## Results

### Genotoxic stress impairs adipogenic competence and induces inflammatory reprogramming in adipocyte progenitors

To establish a model of persistent genotoxic stress, we treated proliferating 3T3-L1 adipocyte progenitors with the topoisomerase II inhibitor etoposide for 48 h before adipogenic induction (**Fig. 1A**). Etoposide produced a dose-dependent loss of adipogenic competence with lipid accumulation at day 8 decreased from 37.7% in vehicle-treated cells to 22% following 0.3 µM etoposide (p = 0.151), 9% following 0.4 µM (p = 0.012) and 5% following 0.5 µM (p = 0.006) (**Fig. 1B,C**).

**Figure 1.**
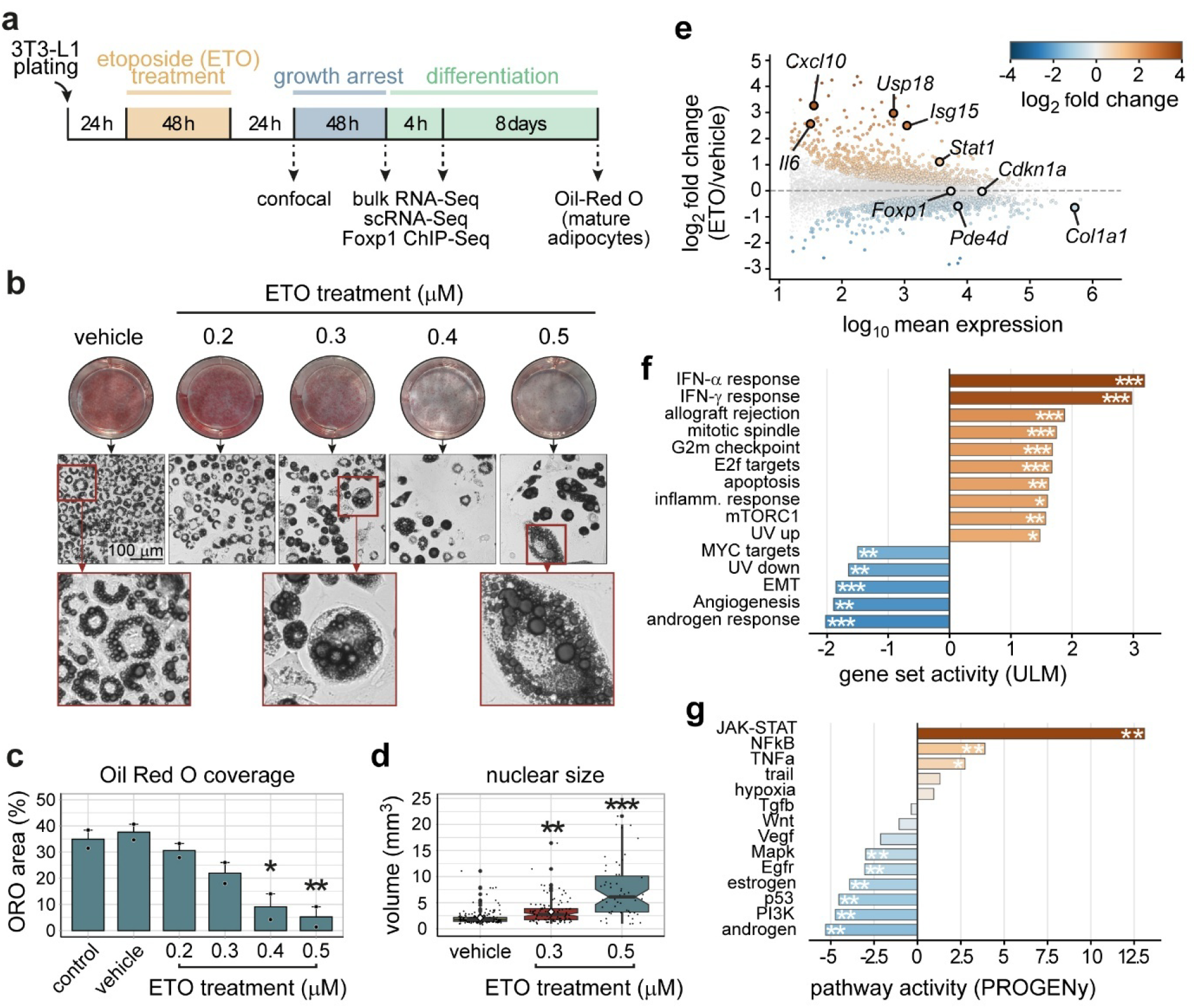
Etoposide induces a senescence-like, adipogenesis-impaired state dominated by a type-I interferon programme. (a) Experimental schematic. 3T3-L1 pre-adipocytes were plated for 24 h, treated with vehicle (DMSO) or etoposide (ETO, 0.2–0.5 µM) for 48 h, allowed 24 h drug-free recovery, growth-arrested for 48 h, then induced with adipogenic differentiation medium. Confocal imaging was performed at the end of growth arrest; bulk RNA-Seq, scRNA-Seq and Foxp1 ChIP-Seq at 4 h post-induction; Oil Red O quantification after 8 days of differentiation. (b) Representative whole-well Oil Red O staining (top), bright-field images with the boxed region indicated in red (middle) and high-magnification insets of lipid droplet morphology (bottom), across vehicle and four etoposide doses. Scale bar 100 µm. (c) Oil Red O area coverage per well (mean ± SEM; n = 2 biological replicates per condition; *p < 0.05, **p < 0.01 versus vehicle, one-way ANOVA with Tukey HSD). (d) Nuclear volume in vehicle, 0.3 µM and 0.5 µM etoposide (boxes, IQR; whiskers, range; individual nuclei overlaid; **p < 0.01, ***p < 0.001 versus vehicle). (e) MA plot of differential expression at 4 h post-induction, etoposide versus vehicle (DESeq2, n = 3 biological replicates per condition), coloured by log₂ fold change. Interferon-stimulated genes (Cxcl10, Usp18, Isg15, Stat1), Il6 and the senescence marker Cdkn1a are induced; Col1a1, Foxp1 and Pde4d are repressed. (f) Gene-set activity by univariate linear modelling across MSigDB Hallmark sets. IFN-α and IFN-γ responses are the most strongly positive; MYC targets, EMT, angiogenesis and androgen response are negative (***p_adj < 0.001, **p_adj < 0.01, *p_adj < 0.05). (g) PROGENy pathway activity. JAK-STAT shows the strongest positive activity, followed by NFκB and TNFα; p53, PI3K, MAPK, EGFR and androgen signalling are negative.

Etoposide-treated cells also developed prominent nuclear abnormalities, including nuclear enlargement, multinucleation and fragmentation (**Fig. 1B,D & Suppl. Fig. 1A**). Nuclear volume increased dose-dependently from 2.0 µm³ in vehicle-treated cells to 3.3 µm³ following 0.3 µM etoposide (p = 0.003) and 7.9 µm³ following 0.5 µM (p < 0.001) (**Fig. 1D**). Nuclear p53 protein increased approximately 3.8-fold following 0.5 µM etoposide (β = 1.341, p = 0.004), whereas Trp53 transcript abundance remained unchanged across several post-treatment timepoints (**Suppl. Fig. 1C-E**), consistent with post-translational stabilisation. Lamin B1 immunofluorescence also increased following genotoxic stress (**Suppl. Fig. 1A-B**). Together, these findings establish a model in which low-dose etoposide produces persistent DNA-damage signalling, loss of adipogenic competence and senescence-associated cellular features.

To determine how this state alters inflammatory transcription, we performed bulk RNA-seq on vehicle- and etoposide-treated progenitors 4 h following adipogenic induction. Differential expression analysis identified 349 genes at |log₂FC| > 1 and adjusted p < 0.05, comprising 273 upregulated and 76 downregulated genes (**Fig. 1E; Suppl. Fig. 2A & Suppl. Table 2**). Among the most strongly induced genes were canonical inflammatory and interferon-responsive genes, including Cxcl10, Usp18, Isg15, Stat1 and Il6 (**Fig. 1E**).

Pathway-level analysis revealed a dominant interferon response. Hallmark IFNα and IFNγ response programmes were the most strongly increased transcriptional signatures, with inferred activities substantially exceeding those of other upregulated programmes (**Fig. 1F & Suppl. Fig. 2B**). PROGENy independently identified JAK-STAT as the most strongly activated signalling pathway, while NFκB and TNFα signalling were also increased (**Fig. 1G**). Complementary transcription-factor (CollecTRI) and STRING network analyses similarly identified interferon-responsive transcription as the dominant response to etoposide (**Suppl. Fig. 2C & 3; Suppl. Table 3**). Thus, persistent genotoxic stress elicits a strong inflammatory transcriptional response dominated by IFN-associated gene expression but accompanied by substantial NFκB-associated signalling.

### Single-cell analysis identifies Foxp1 and Pde4d expression as associated with distinct IFN and NFκB-associated damage states

The simultaneous engagement of IFN- and NFκB-associated programmes in bulk transcriptomes raised an important question: are these responses uniformly co-engaged within damaged cells, or do they reflect heterogeneity in inflammatory state? To resolve the cellular organisation of the response, we performed Parse-seq across vehicle- and etoposide-treated 3T3-L1 progenitors before and shortly after adipogenic induction. Following quality filtering, 3,966 cells were retained and partitioned into transcriptionally distinct populations (**Fig. 2A & Suppl. Fig. 4**). Pseudobulk changes closely reproduced the bulk RNA-seq response (r > 0.9 across STRING-derived transcriptional modules; **Suppl. Fig. 4D**), confirming that the single-cell dataset captured the major consequences of genotoxic stress.

**Figure 2.**
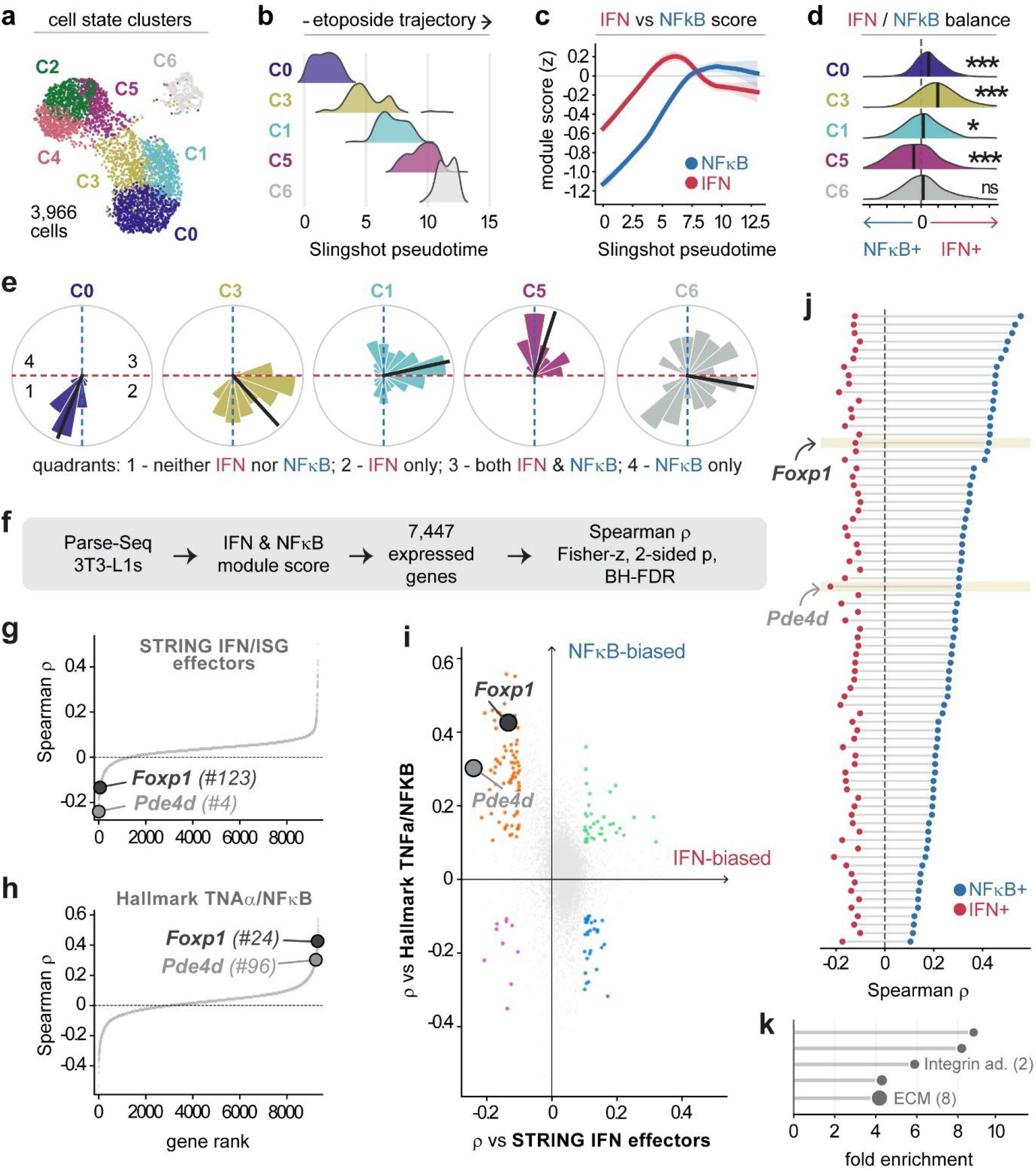
Single-cell analysis resolves inflammatory configuration, and a per-cell screen identifies Foxp1 and Pde4d as anti-IFN, pro-NFκB candidates. (a) UMAP of 3,966 Parse-seq 3T3-L1 cells across four conditions (vehicle or etoposide, each at 0 h and 4 h post-induction), coloured by the seven cell-state clusters C0–C6. (b) Ridge plot of Slingshot pseudotime along the etoposide damage trajectory for the clusters lying on that axis (C0, C3, C1, C5, C6). (c) IFN (red) and NFκB (blue) module scores as a function of Slingshot pseudotime (z-scored, loess fit with 95% CI). The IFN programme peaks mid-trajectory while NFκB rises later and is sustained. (d) Distribution of the per-cell IFN/NFκB balance by cluster (negative, NFκB-biased; positive, IFN-biased). Black bars, cluster medians. Asterisks, one-versus-rest test with BH correction (***q < 0.001, *q < 0.05; ns, not significant). (e) Polar histograms of per-cell IFN/NFκB configuration by cluster. Quadrants: 1, neither programme; 2, IFN only; 3, both; 4, NFκB only. Black line, circular mean. Clusters differ in which configuration they occupy rather than in overall inflammatory magnitude. (f) Screen design. For each of 7,447 expressed genes, per-cell Spearman correlations were computed against an IFN and an NFκB module score, with Fisher-z transformation, two-sided p-values and BH-FDR correction. (g) Waterfall of gene rank versus Spearman ρ for the STRING IFN/ISG effector module; Pde4d is the 4th most anti-correlated gene genome-wide. (h) Equivalent waterfall for Hallmark TNFα/NFκB; Foxp1 ranks 24th and Pde4d 96th, both positively correlated. (i) Per-gene ρ versus the STRING IFN effector module (x) against ρ versus Hallmark TNFα/NFκB (y). Genes meeting an effect-size threshold of |ρ| ≥ 0.10 on both axes are coloured by quadrant; the anti-IFN, pro-NFκB quadrant is shown in orange. Foxp1 and Pde4d sit deep within it. (j) Paired per-gene correlations for the anti-IFN, pro-NFκB quadrant genes. Each row is one gene, showing ρ versus IFN (red) and ρ versus NFκB (blue) joined by a grey line, ranked by NFκB correlation. Foxp1 and Pde4d are highlighted (shaded rows). (k) Functional over-representation among the quadrant genes, shown as fold enrichment against the 7,447 screened genes as background. The set is dominated by extracellular matrix and adhesion terms rather than by inflammatory annotation.

Trajectory inference ordered the principal etoposide-responsive populations along a transcriptional axis extending from baseline progenitors through distinct damage-associated states (**Fig. 2B & Suppl. Fig. 4E**). IFN-associated transcription increased rapidly along this trajectory, reached a maximum at intermediate pseudotime and subsequently declined, whereas NFκB-associated transcription increased more gradually and remained elevated later in the trajectory (**Fig. 2C**). Consequently, cell populations differed substantially in their relative engagement of the two programmes: early damage-associated states were predominantly IFN-biased, whereas later states showed relatively greater NFκB-associated transcription (**Fig. 2D,E**). This ordering is consistent with a transition from an early IFN-dominant response towards greater NFκB engagement, although pseudotime represents transcriptional ordering rather than direct longitudinal tracking of individual cells.

We next exploited this heterogeneity to identify genes associated with inflammatory-state configuration. For each of 7,447 detected genes, we calculated its per-cell relationship with independently defined IFN/ISG and TNFα/NFκB transcriptional scores (**Fig. 2F**). As expected, genes positively associated with the IFN score were dominated by canonical interferon-stimulated genes (**Suppl. Table 4**). By contrast, Pde4d, encoding a major cAMP phosphodiesterase, ranked fourth genome-wide among genes negatively associated with IFN-associated transcription (ρ = −0.160, adjusted p = 8.4 × 10⁻²²), while Foxp1 ranked within the top 1.7% (ρ = −0.097, adjusted p = 3.5 × 10⁻⁸) (**Fig. 2G**).

Strikingly, both genes showed the opposite relationship with NFκB-associated transcription. Foxp1 ranked 24th genome-wide (ρ = +0.42) and Pde4d 96th (ρ = +0.30) among genes positively associated with the TNFα/NFκB score (**Fig. 2H**). Across the transcriptome, Foxp1 and Pde4d therefore occupied a relatively restricted sector characterised by lower IFN- and higher NFκB-associated transcription (**Fig. 2I,J**). We then asked what kind of gene occupies this sector. Over-representation analysis of the 75 genes meeting an effect-size threshold on both axes returned only structural terms: extracellular matrix components were enriched 10.3-fold over the screened background, comprising five collagens together with Vcan, Sparc, Sned1 and two ADAMTS proteases, and adhesion and cytoskeletal regulators were enriched 23.5-fold (**Fig. 2K; Suppl. Table S5**). No interferon, NFκB or cytokine term reached significance. The set was not devoid of immune genes, but those it contained were receiving rather than effector components: Il1r1, Il1rl1 and the IL-1 receptor-associated adaptor Peli2, together enriched 29.8-fold, with no chemokine or cytokine genes represented. Foxp1 and Pde4d therefore associate with a transcriptional configuration defined largely by matrix production and adhesion, in cells equipped to receive IL-1 family input. Together, these analyses identify Foxp1 and Pde4d independently through inflammatory-state heterogeneity and raise the possibility that they form a functionally related regulatory pair.

### Foxp1 positively regulates Pde4d and inflammatory transcriptional state

The concordant association of Foxp1 and Pde4d with inflammatory state raised the possibility that the two genes are mechanistically linked. We therefore examined Foxp1 chromatin occupancy in vehicle- and etoposide-treated 3T3-L1 progenitors. Genome-wide Foxp1 occupancy remained broadly stable following etoposide treatment, but differential-binding analysis identified five reproducible damage-responsive sites at FDR < 0.05 (**Fig. 3A,B, Suppl. Figs. 5 & 6; Suppl. Table 6**). Foxp1 occupancy was lost on etoposide at sites associated with Pde4d, Specc1 and Jag1, and gained at sites associated with Acp7 and Tek (**Fig. 3A,B**). The strongest loss of Foxp1 occupancy occurred immediately 3′ of Pde4d (**Fig. 3A-C & Suppl. Fig. 6A**).

**Figure 3.**
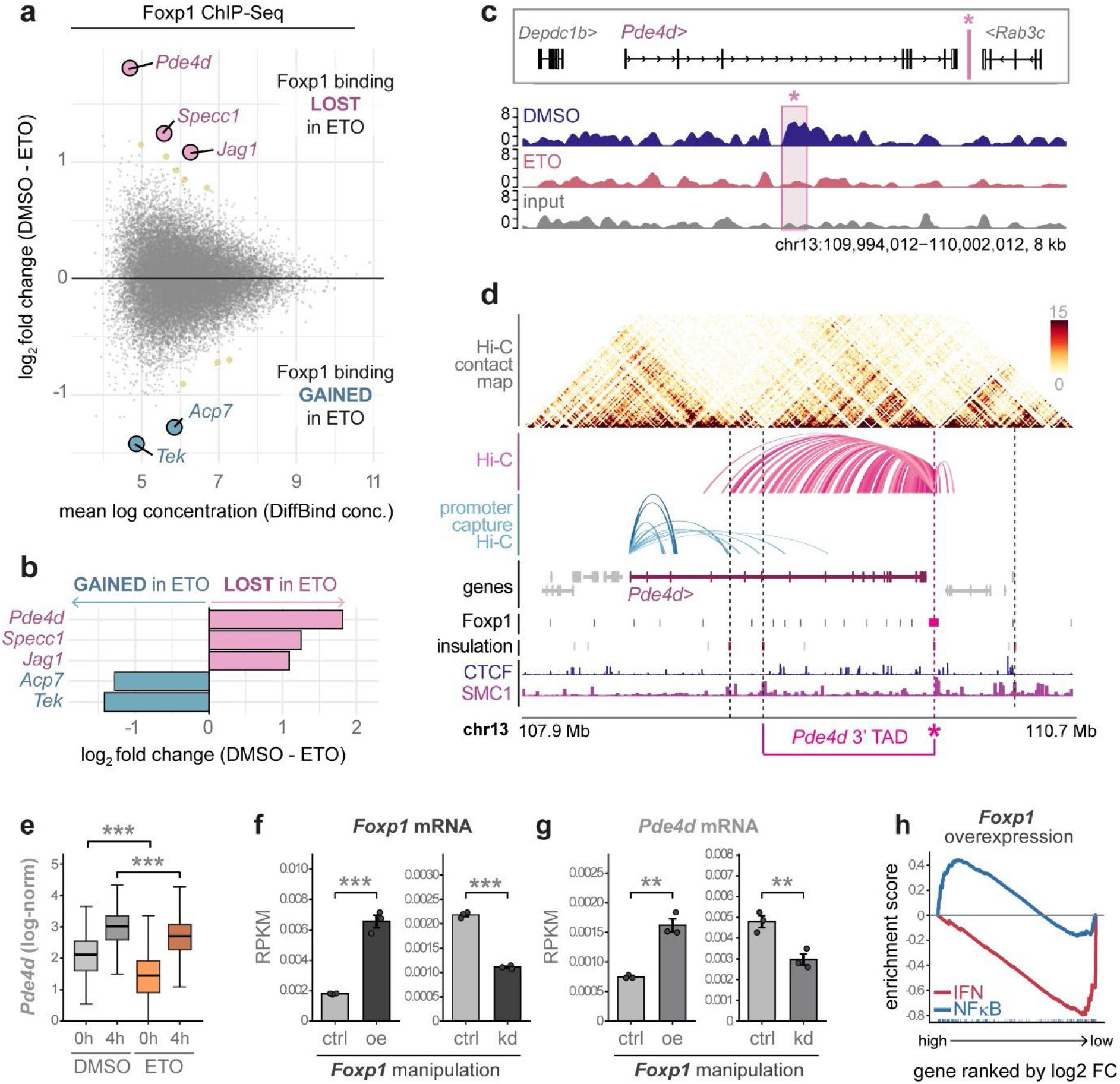
Foxp1 occupies a damage-sensitive regulatory element within the Pde4d 3′ TAD and positively regulates Pde4d. (a) MA plot of differential Foxp1 binding (DiffBind v3) in etoposide-versus vehicle-treated 3T3-L1 cells at 4 h post-induction. Positive values indicate binding lost on etoposide. Five peaks reach significance: Pde4d, Specc1 and Jag1 lost; Acp7 and Tek gained. (b) Log₂ fold change (DMSO − ETO) for the five differentially bound peaks. (c) Foxp1 ChIP-Seq coverage at the Pde4d 3′ element (chr13:109,994,012–110,002,012, 8 kb) in DMSO (blue), etoposide (red) and input (grey). The element (asterisk, shaded) lies downstream of Pde4d and upstream of Rab3c. Tracks are RPGC-normalised. (d) Chromatin architecture across chr13:107.9–110.7 Mb. From top: Hi-C contact map; Hi-C arcs and promoter-capture Hi-C arcs from the element; gene models; Foxp1 peaks, with the damage-sensitive element in magenta; insulation score; CTCF and SMC1 occupancy. The element lies at the 3′ boundary of the Pde4d TAD. (e) Pde4d expression (log-normalised) in vehicle and etoposide at 0 h and 4 h post-induction (***p < 0.001). (f) Foxp1 mRNA (RPKM) following Foxp1 overexpression (oe) and knockdown (kd) versus matched controls in 3T3-L1 (GSE179031; ***p < 0.001). (g) Pde4d mRNA (RPKM) in the same samples, showing reciprocal change in both directions (**p < 0.01). (h) Gene-set enrichment on genes ranked by log₂ fold change following Foxp1 overexpression. The STRING IFN/ISG effector module is depleted (red) while Hallmark TNFα/NFκB is enriched (blue).

This damage-sensitive Foxp1-bound region was located within a Pde4d chromatin domain close to a chromatin structural boundary (**Fig. 3D & Suppl. Fig. 6A**). Integration with high-resolution Hi-C data showed 66.9% of 2,521 interactions originating from the Foxp1-bound region terminate within the Pde4d gene body (**Fig. 3D & Suppl. Fig. 6A**). With Hi-C interactions preferentially contacting short Pde4d isoforms (**Suppl. Fig. 7 & Suppl. Table 7**). These observations link the damage-sensitive Foxp1-binding site to Pde4d.

Consistent with reduced Foxp1 occupancy, Pde4d expression decreased following etoposide treatment (**Fig. 3E**). We therefore re-analysed independent RNA-seq datasets in which Foxp1 abundance had been experimentally manipulated in 3T3-L1 cells (Ma et al. 2023). Foxp1 overexpression increased Pde4d expression approximately two-fold, whereas Foxp1 knockdown reduced Pde4d by approximately 38% (**Fig. 3F,G**). Thus, Foxp1 positively regulates Pde4d expression, providing a mechanistic explanation for their concordant behaviour in the single-cell dataset.

We next asked whether experimentally altering Foxp1 abundance was sufficient to modify the inflammatory transcriptional programmes through which Foxp1 had been identified. Foxp1 overexpression suppressed IFN-associated transcription while increasing TNFα/NFκB-associated transcription (**Fig. 3H**), reproducing the direction of the relationships observed across individual damaged cells. Together, these findings identify Pde4d as a positive Foxp1 transcriptional target and link Foxp1 abundance to the relative configuration of IFN- and NFκB-associated transcriptional programmes.

### Foxp1-Pde4d coupling and its association with inflammatory state are conserved across vertebrate systems

Having established a regulatory relationship between Foxp1 and Pde4d in 3T3-L1 cells, we next asked whether their coupling extends across biological contexts. We analysed Foxp1-Pde4d co-expression across 82 tissues from mouse, human and turquoise killifish, spanning approximately 430 million years of vertebrate evolution (**Fig. 4A & Suppl. Table 8**). Foxp1 and Pde4d expression were positively correlated in 79 of 82 tissues (96%), with significant positive associations after multiple-testing correction in 72 tissues (88%) (**Fig. 4A,B**).

**Figure 4.**
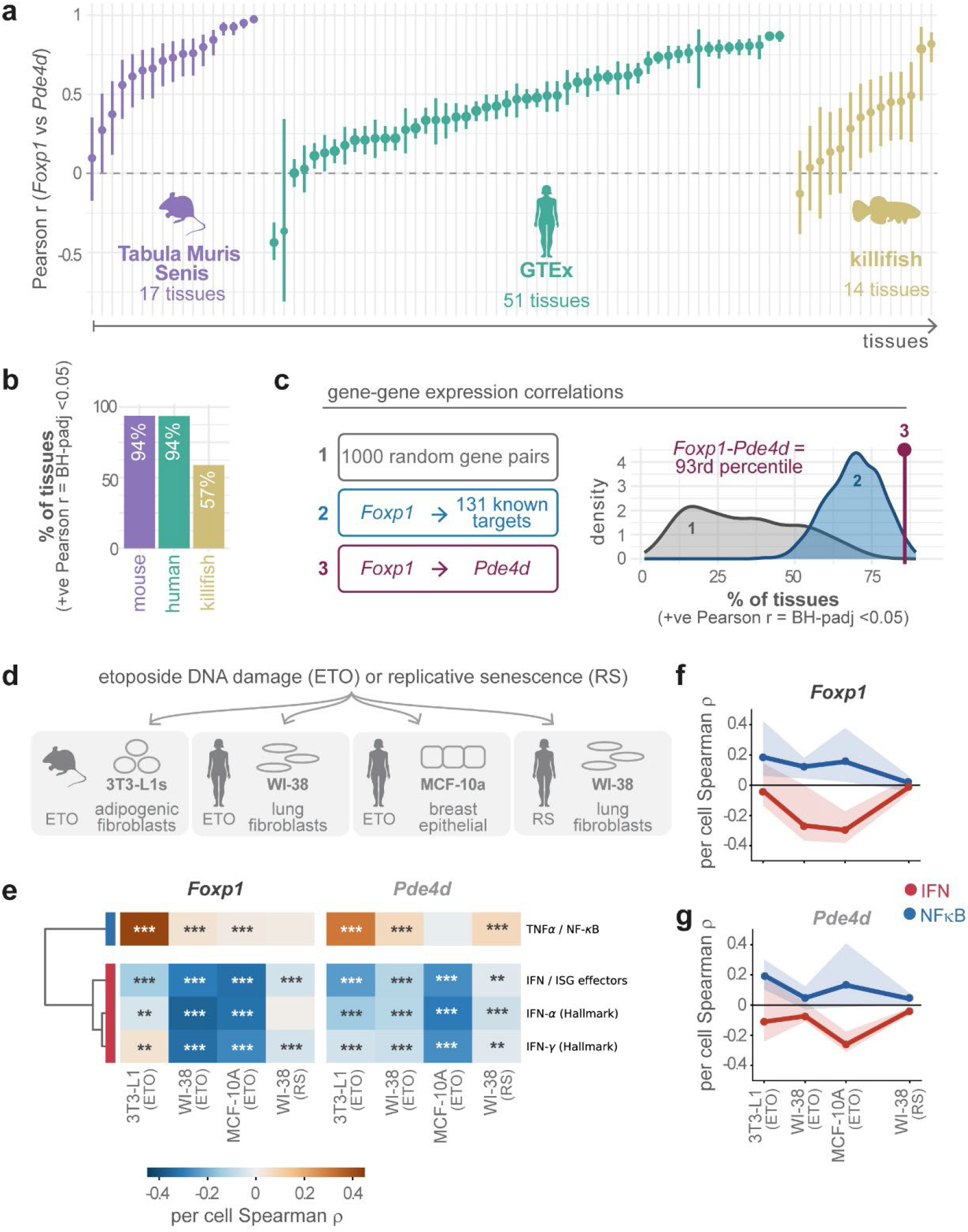
The Foxp1-Pde4d relationship is conserved across vertebrates and reproduces across independent damage and senescence systems. (a) Within-tissue Pearson correlation between Foxp1 and Pde4d across tissues of three vertebrate atlases: mouse Tabula Muris Senis (17 tissues), human GTEx v8 (51 tissues) and turquoise killifish (14 tissues). Points, correlation; bars, 95% CI; tissues ordered by correlation within species. (b) Percentage of tissues per species with a significantly positive correlation (BH-adjusted p < 0.05). (c) Conservation benchmark. The Foxp1–Pde4d conservation index is compared against 1,000 random gene pairs (grey) and 131 high-confidence Foxp1-activated target genes (blue), scored under identical per-tissue thresholds. Foxp1–Pde4d falls at the 93rd percentile. (d) Four single-cell cohorts spanning etoposide-induced DNA damage (ETO) and replicative senescence (RS): 3T3-L1 adipogenic fibroblasts, WI-38 lung fibroblasts, MCF-10A breast epithelium, and replicatively senescent WI-38. (e) Per-cell Spearman correlations of Foxp1 and Pde4d against four inflammatory modules in each cohort. The negative association with the three interferon modules is reproduced in every cohort; the positive association with TNFα/NFκB is present in most but not all (***q < 0.001, **q < 0.01). (f, g) The same correlations for Foxp1 (f) and Pde4d (g) shown across cohorts, with IFN in red and NFκB in blue. Shaded bands, 95% CI.

To determine whether this degree of conservation exceeded that expected for arbitrary gene pairs, we compared Foxp1-Pde4d with 1,000 randomly sampled pairs and a curated set of 131 Foxp1-activated targets. Foxp1-Pde4d ranked among the most consistently positively coupled relationships across tissues and ranked in the 93^rd^ percentile compared to known Foxp1-target pairs (**Fig. 4C**). Foxp1-Pde4d were also positively correlated at the single-cell level in human adipose tissue, and their expression levels co-declined with age (**Suppl. Fig. 8**). Thus, the relationship identified in adipocyte progenitors represents an unusually conserved transcriptional association across divergent vertebrate lineages and tissue contexts.

We next asked whether the Foxp1-Pde4d relationship with inflammatory state similarly generalised beyond the 3T3-L1 discovery system. We analysed three independent human single-cell datasets encompassing etoposide-treated WI-38 fibroblasts, etoposide-treated MCF-10A epithelial cells and replicative senescent WI-38 fibroblasts, together comprising more than 60,000 cells with our 3T3-L1 dataset (**Fig. 4D & Suppl. Fig. 9**).

The inverse relationship with IFN-associated transcription was highly reproducible. Foxp1 expression was negatively associated with IFN/ISG transcription across all four datasets (ρ = −0.13 to −0.33), as was Pde4d (ρ = −0.03 to −0.27) (**Fig. 4E-G**). Positive relationships with NFκB-associated transcription were also directionally reproduced but varied more substantially between systems, being strongest during acute damage and weak during replicative senescence (**Fig. 4E-G**). Thus, Foxp1-Pde4d coupling is broadly conserved across vertebrate tissues, and its relationship with inflammatory state is also conserved, with the inverse association with IFN representing the most robust feature across damage and senescence models. Consistent with conservation of this relationship in vivo, analysis of an independent zebrafish fin-regeneration dataset identified coordinated *foxp1b*-*pde4d* expression during regeneration, with coupling concentrated within intermediate epithelial states (**Suppl. Fig. 12**).

### Foxp1 is required for inducible macrophage recruitment following tissue injury

Having identified Foxp1-Pde4d through variation in inflammatory state, we next asked whether Foxp1 also influences the capacity to respond to tissue challenge in vivo. We used zebrafish, in which mammalian Foxp1 is represented by the co-orthologues *foxp1a* and *foxp1b*, and generated F0 larvae with CRISPR-mediated disruption of both genes (*foxp1ab*) (**Fig. 5A**) (Wafer et al. 2026). We performed tail-fin injury which was followed by longitudinal imaging of neutrophil (*lyzC:DsRed*) and macrophage (*mfap4:mTurquoise*) reporters, with immune-cell responses quantified within a 500-µm region surrounding the wound (**Fig. 5A,B**).

**Figure 5.**
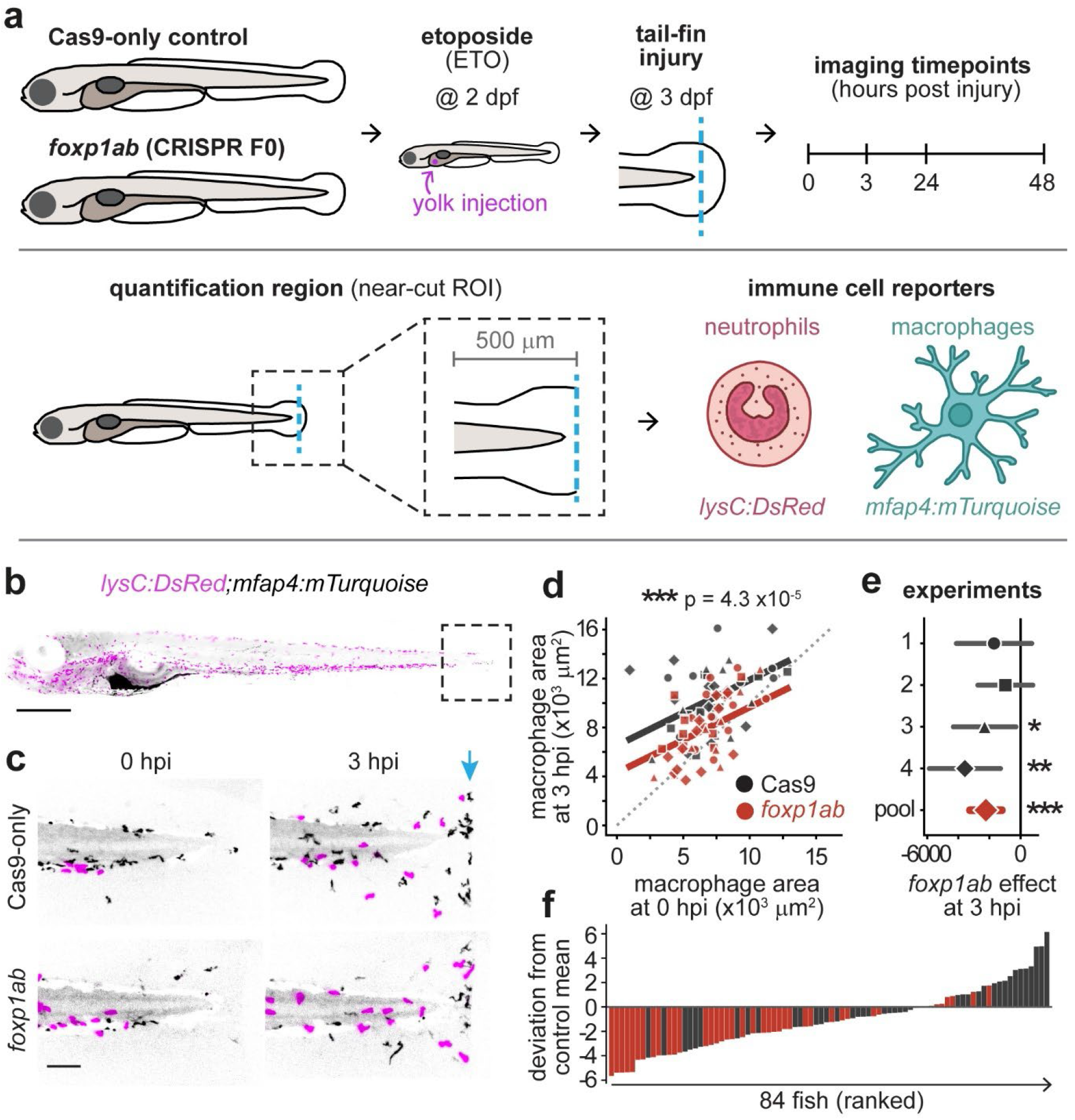
Foxp1 is required for inducible macrophage recruitment following tail-fin injury. (a) Experimental design. Embryos were injected at the one-cell stage with *foxp1a* and *foxp1b* guide RNPs (*foxp1ab* crispants) or with Cas9 alone, yolk-injected with etoposide or vehicle at 2 dpf, tail-fin transected at 3 dpf, and imaged at 0, 3, 24 and 48 hpi. Leukocytes were quantified within a 500 µm region of interest proximal to the cut, using *Tg(lysC:dsRed2)* for neutrophils and *Tg(mfap4.1:mTurquoise)* for macrophages. (b) Representative whole larva carrying both reporters, with the imaged tail region boxed. Scale bar 250 µm. (c) Representative near-cut projections at 0 and 3 hpi in Cas9-only and *foxp1ab* larvae. Macrophage signal, black; neutrophil signal, magenta. Blue arrow, transection plane. Scale bar 100 µm. (d) Macrophage area within the near-cut region at 3 hpi against the same fish’s 0 hpi value, for Cas9-only (black) and *foxp1ab* (red). Symbols denote experiment; lines, fitted ANCOVA; dotted line, unity. The genotype effect conditioning on baseline is p = 4.3 × 10⁻⁵. (e) *foxp1ab* effect on macrophage area at 3 hpi estimated within each of the four experiments and pooled. Points, estimate; bars, 95% CI (*p < 0.05, **p < 0.01, ***p < 0.001). (f) Per-fish deviation from the control mean, all 84 fish ranked. Grey, Cas9-only; red, *foxp1ab*. The deficit is a shift of the whole distribution rather than a subset of outliers.

The clearest phenotype was observed in macrophages. Prior to injury, macrophage abundance was comparable between Cas9 controls and *foxp1ab* larvae (ratio 0.93, p = 0.30). Following injury, however, macrophage accumulation was markedly reduced in *foxp1ab* animals (**Fig. 5C**). Conditioning 3-h post-injury macrophage area on the corresponding pre-injury value from the same animal identified a significant Foxp1-dependent recruitment deficit across four independent experiments (pooled effect −2,224 µm²; −21.5%; p < 0.0001; n = 84) (**Fig. 5D,E & Suppl. Table 9**). The phenotype was evident across individual animals rather than being driven by a small number of outliers (**Fig. 5F**). A similar, but smaller, reduction in neutrophil recruitment to the injury at 3 hpi was also observed in *foxp1ab* larvae (p = 0.018) (**Fig. 5C & Suppl. Fig. 11**). The Foxp1-dependent recruitment deficit was evident both with and without prior etoposide exposure, with no genotype-by-etoposide interaction, indicating that impaired responsiveness did not require experimentally imposed genotoxic stress (**Suppl. Fig. 11**).

Thus, Foxp1 deficiency does not substantially alter macrophage abundance before injury but markedly impairs their subsequent recruitment to damaged tissue. Foxp1 is therefore required for effective inducible macrophage recruitment following tissue challenge.

### Foxp1 preserves the inducible range of NFκB signalling through a cAMP-sensitive mechanism

Reduced macrophage recruitment could reflect a weaker injury cue, or a loss of inducibility imposed by an altered resting state. To distinguish these possibilities, we monitored NFκB activity longitudinally following tail-fin injury using an NFκB:eGFP reporter (**Fig. 6A**) (Kanther et al. 2011).

**Figure 6.**
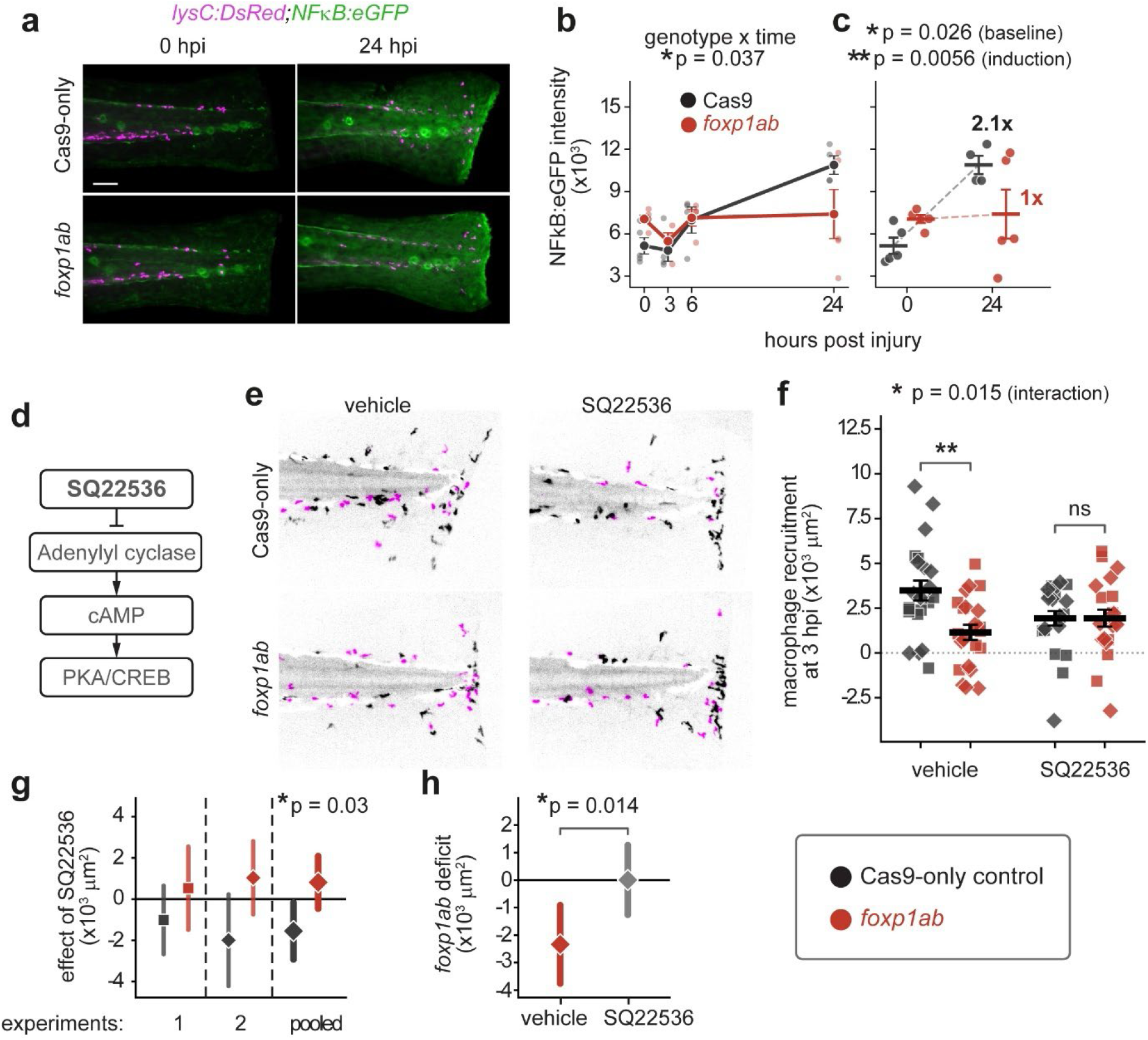
Foxp1 preserves the inducible range of NFκB signalling through a cAMP-sensitive mechanism. (a) Representative near-cut projections of larvae carrying Tg(lysC:dsRed2) (magenta) and an NFκB:eGFP reporter (green) at 0 and 24 hpi in Cas9-only and foxp1ab animals. Scale bar 100 µm. (b) Near-cut NFκB:eGFP intensity at 0, 3, 6 and 24 hpi in Cas9-only (black) and foxp1ab (red). Points, individual larvae; line, group mean ± SEM. Genotype by time interaction, p = 0.037. (c) The same data at 0 and 24 hpi. foxp1ab larvae have higher pre-injury reporter activity (baseline, p = 0.026) and lose the injury-induced increase seen in controls (induction, p = 0.0056); fold induction 2.1× in Cas9-only versus 1× in foxp1ab. (d) SQ22536 inhibits adenylyl cyclase, reducing cAMP production and downstream PKA/CREB signalling. (e) Representative near-cut projections at 3 hpi in vehicle and SQ22536, for Cas9-only and *foxp1ab* larvae. Macrophage signal, black; neutrophil signal, magenta. (f) Macrophage recruitment at 3 hpi, baseline-adjusted, by genotype and treatment. Symbols denote experiment; bars, mean ± SEM. Genotype by treatment interaction, p = 0.015; the genotype difference is significant under vehicle (**) and absent under SQ22536 (ns). (g) Effect of SQ22536 within each genotype, estimated separately in the two contributing experiments and pooled (p = 0.03). Grey, Cas9-only; red, *foxp1ab*. The drug raises recruitment in mutants and lowers it in controls. (h) *foxp1ab* deficit measured separately within each treatment arm, showing that the deficit present under vehicle is abolished under SQ22536 (p = 0.014). Points, estimate; bars, 95% CI.

Before injury, NFκB reporter activity was significantly elevated in *foxp1ab* larvae relative to controls (p = 0.026), demonstrating that Foxp1 loss does not produce a constitutively hypo-inflammatory state (**Fig. 6B,C**). The major difference instead emerged following injury. NFκB reporter activity increased approximately 2.1-fold in controls but showed essentially no net induction in *foxp1ab* animals (p = 0.0056 for induction), producing a significant genotype-by-time interaction across the response (p = 0.037) (**Fig. 6B,C**). Thus, elevated basal NFκB activity following Foxp1 loss is accompanied by markedly reduced capacity for further activation following tissue damage, compressing the inducible range of NFκB signalling.

Because Pde4d terminates cAMP signalling, we hypothesised that altered cAMP regulation contributes to the impaired inflammatory response following Foxp1 loss. We therefore pre-treated larvae with the adenylyl cyclase inhibitor SQ22536 - which reduces cAMP generation – from two-hours before tail-fin transection and quantified macrophage recruitment following injury (**Fig. 6D,E**). In vehicle-treated animals, *foxp1ab* larvae again showed substantially reduced macrophage recruitment relative to controls. SQ22536 shifted recruitment in opposite directions in the two genotypes, reducing recruitment in controls while increasing it in *foxp1ab* larvae and thus eliminating the recruitment deficit (genotype-by-treatment interaction p = 0.015; **Fig. 6F**).

This genotype-dependent effect was reproduced across two independent experiments and was significant when the experiments were pooled (interaction p = 0.014; n = 87) (**Fig. 6G,H**). Adenylyl cyclase inhibition therefore restores the principal functional consequence of Foxp1 loss. Together with the positive regulation of Pde4d by Foxp1 in adipocyte progenitors, these findings implicate altered cAMP signalling in the impaired inflammatory responsiveness of Foxp1-deficient animals.

Together, our findings support a model in which Foxp1-dependent Pde4d expression contributes to a cAMP signalling environment that preserves inflammatory responsiveness (**Fig. 7**). In the intact state, Foxp1 positively regulates Pde4d, supporting controlled cAMP signalling, relatively low basal NFκB activity and preserved capacity for NFκB induction following tissue injury. Foxp1 deficiency reduces Pde4d expression and alters cAMP signalling, a state associated in vivo with elevated basal NFκB activity, reduced injury-induced NFκB activation and impaired macrophage recruitment. Pharmacological rescue by adenylyl cyclase inhibition further implicates cAMP-sensitive signalling in this phenotype. Thus, Foxp1 loss does not simply increase or decrease inflammatory signalling, but alters the range over which the inflammatory response can adapt to tissue challenge.

**Figure 7.**
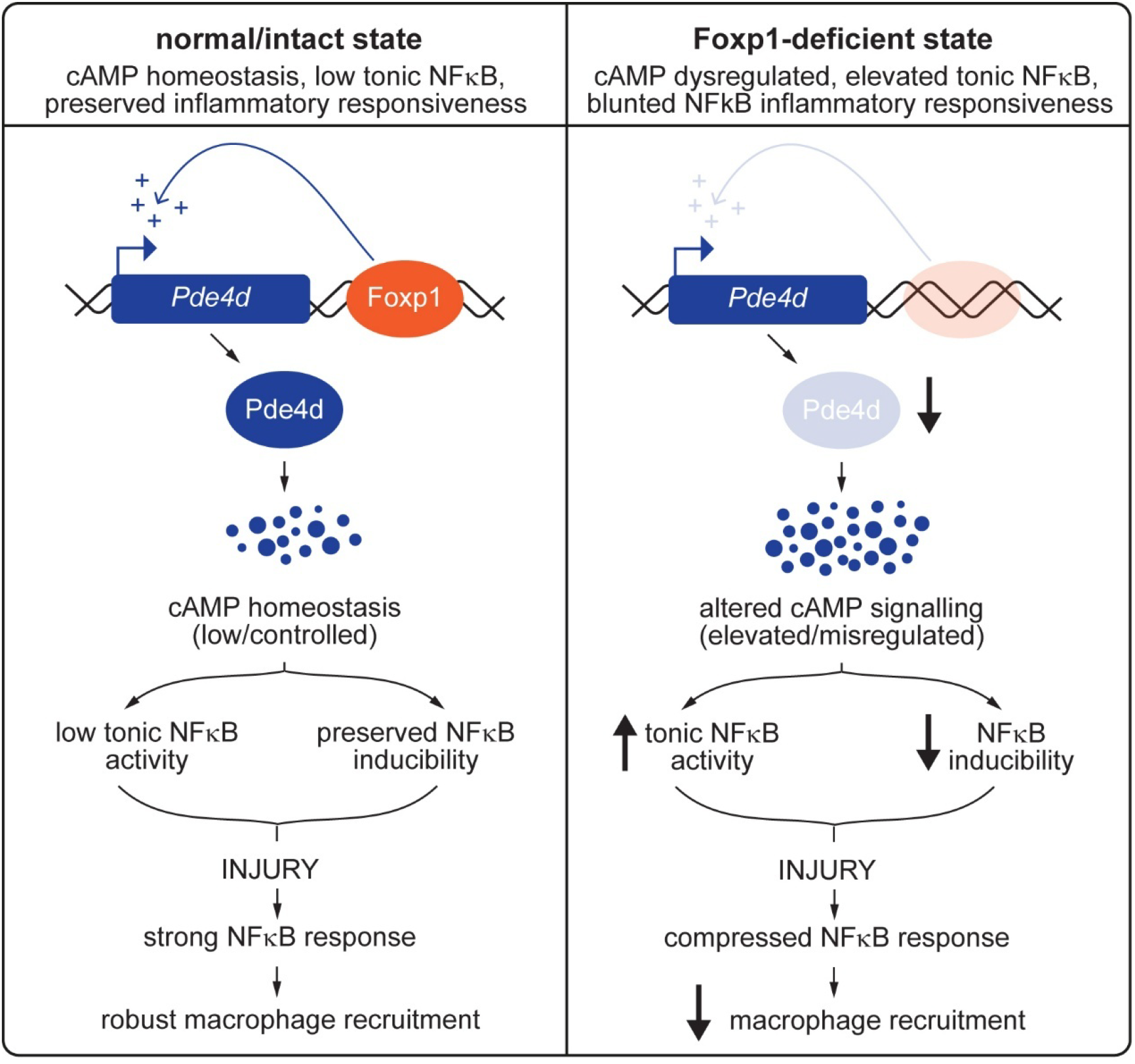
Model: Foxp1-dependent Pde4d expression sets the cAMP environment from which an inflammatory response is mounted. Left, intact state. Foxp1 occupies a regulatory element at the Pde4d locus and positively supports Pde4d expression. Pde4d-dependent cAMP hydrolysis maintains cAMP homeostasis, which is associated with low tonic NFκB activity and preserved NFκB inducibility, so injury produces a strong NFκB response and robust macrophage recruitment. Right, Foxp1-deficient state. Loss of Foxp1 reduces Pde4d expression and alters cAMP signalling, which is associated with elevated tonic NFκB activity and reduced NFκB inducibility, so the same injury produces a compressed NFκB response and impaired macrophage recruitment. Arrow weight indicates the direction and relative magnitude of change.

## Discussion

Inflammatory responses to cellular damage are heterogeneous, reflecting differences in cell identity, stimulus, signalling history and time. Here, we used this heterogeneity to identify a conserved regulatory relationship linking cellular state to inflammatory responsiveness. Our principal findings are that: (i) genotoxic stress generates distinct configurations of IFN- and NFκB-associated transcription within adipocyte progenitors; (ii) Foxp1 and Pde4d emerge from unbiased analysis as strongly associated with this inflammatory configuration, and their expression is coupled across diverse vertebrate tissues and experimental systems; (iii) Foxp1 positively regulates Pde4d and exhibits damage-sensitive occupancy at a Pde4d-associated regulatory element; and (iv) Foxp1 loss in vivo elevates basal NFκB activity but reduces its subsequent induction and macrophage recruitment following injury, with recruitment restored by adenylyl cyclase inhibition. Together, these findings identify Foxp1-Pde4d as a previously unrecognised regulatory relationship connecting cellular state, cAMP regulation and the capacity to mount an inflammatory response.

A central feature of our study is that inflammatory-state heterogeneity provided the means to identify Foxp1-Pde4d. Building on evidence that cell-to-cell variation following genotoxic stress reflects structured biological states rather than stochastic noise (Burnaevskiy et al. 2023; Wechter et al. 2023; Jang et al. 2026), we resolved IFN- and NFκB-associated transcription independently and used their relative engagement as a discovery space for candidate regulators. This revealed variation not simply in inflammatory magnitude, but in inflammatory configuration. Foxp1 and Pde4d independently occupied a relatively restricted IFN-low/NFκB-high region of this transcriptional space and were subsequently found to be functionally linked. This relationship extended well beyond the system in which it was discovered: Foxp1 and Pde4d expression was positively coupled across 79 of 82 vertebrate tissues spanning approximately 430 million years of evolution, while their inverse relationship with IFN- and positive relationship with NFκB-associated transcription was reproduced across independent models of genotoxic stress and senescence. Thus, exploiting inflammatory heterogeneity exposed a regulatory relationship that generalised across species, tissues and experimental contexts.

The emergence of Foxp1 and Pde4d as a linked pair is particularly interesting in light of recent evidence connecting Foxp1 to cyclic-nucleotide phosphodiesterase regulation in another biological context. Foxp1 is a broadly expressed transcription factor with established roles in development and cell-state specification, and in humans FOXP1 haploinsufficiency causes FOXP1 syndrome, a neurodevelopmental disorder characterised by intellectual disability, speech and language impairment and behavioural abnormalities (Stewart and Cho 2025; Lozano et al. 2021). Recent work in a mouse model of FOXP1 syndrome identified marked loss of the striatal phosphodiesterase Pde10a following Foxp1 haploinsufficiency, accompanied by altered inflammatory and microglial phenotypes (Fröhlich et al. 2025). Our identification of Pde4d as a positive Foxp1 transcriptional target in adipocyte progenitors therefore raises the possibility that regulation of cyclic-nucleotide phosphodiesterases represents a broader, context-dependent output of Foxp1. Unlike the striatally restricted Pde10a relationship, however, Foxp1-Pde4d expression coupling was remarkably widespread, occurring across diverse tissues and vertebrate species. PDE4D regulates the abundance and spatial organisation of cAMP, providing a route through which Foxp1-dependent transcription could influence the cellular signalling environment. The consistency of Foxp1-Pde4d coupling across contexts, rather than exceptionally strong correlation within individual tissues, suggests a recurrent quantitative relationship that is deployed across distinct cellular states rather than a lineage-restricted transcriptional programme.

The cellular state associated with Foxp1-Pde4d provides a further clue to its biological context. Genes sharing the IFN-low/NFκB-high configuration with Foxp1 and Pde4d were enriched not for inflammatory effector programmes, but for extracellular-matrix, adhesion and cytoskeletal programmes. This is consistent with an emerging view of damage and senescence in which inflammatory and tissue-remodelling programmes represent interconnected but separable dimensions of cellular state. Indeed, single-cell studies have identified ECM-remodelling states during DNA-damage-induced senescence and across damaged tissues, suggesting that matrix regulation is an important component of the heterogeneous response to cellular stress (Jang et al. 2026; Evans et al. 2023; Mavrogonatou et al. 2023). In our data, this relationship extended beyond adipocyte progenitors. During zebrafish fin regeneration, *foxp1b*-*pde4d* coupling was concentrated within intermediate epithelial states, including an adhesion-associated state, and was absent from a chemokine-producing epithelial state (**Suppl. Fig. 12**). Interestingly, a related association is evident in human adipose tissue, where PDE4D marks a subpopulation within ADIPOMATlo adipocytes, a non-canonical adipocyte state characterised by inflammatory, pro-fibrotic and extracellular-matrix-associated features (Efthymiou et al. 2026). Together, these observations raise the possibility that Foxp1-Pde4d coupling is preferentially engaged within structural or tissue-remodelling cellular states across otherwise distinct cell types. Rather than representing an isolated inflammatory regulator, Foxp1-Pde4d may therefore form part of a broader state programme that links matrix interaction and cellular architecture to the configuration of inflammatory signalling.

Our data provide a transcriptional mechanism linking Foxp1 to Pde4d that may be particularly consequential given Pde4d isoform architecture. Genome-wide Foxp1 occupancy was largely stable following genotoxic stress, yet ChIP-seq identified selective loss of Foxp1 binding at an active regulatory element 3′ of Pde4d. Together with chromatin contacts across the Pde4d locus and reciprocal changes in Pde4d expression following Foxp1 gain- and loss-of-function, these findings identify Pde4d as a positive Foxp1 transcriptional target and link its regulation to damage-sensitive Foxp1 occupancy. Notably, short Pde4d isoforms predominated in adipocyte progenitors and were preferentially reduced following etoposide treatment. These isoforms lack UCR1 and therefore the canonical PKA-dependent feedback through which rising cAMP can increase PDE4 activity (Baillie et al. 2019; Fertig and Baillie 2018; Kyurkchieva and Baillie 2023; Houslay and Adams 2003), potentially making cAMP regulation more dependent on Pde4d abundance, localisation and alternative regulatory inputs. Thus, relatively slow changes in Foxp1-dependent Pde4d transcription could alter the cAMP signalling environment from which subsequent stimuli are interpreted. Although the function of the 3′ regulatory element, isoform-specific protein abundance and cAMP dynamics remain to be established, these findings provide a potential mechanistic bridge between damage-induced transcriptional state and subsequent signalling responsiveness.

The most striking functional consequence of Foxp1 loss was not a simple increase or decrease in inflammation, but a dissociation between basal inflammatory state and response capacity. In zebrafish, Foxp1 deficiency increased basal NFκB reporter activity yet markedly blunted its subsequent induction following tissue injury, coincident with impaired macrophage recruitment. Thus, greater inflammatory activity at baseline was associated with a reduced capacity to increase that activity when challenged. Elevated basal inflammatory signalling can coexist with impaired responsiveness in settings including inflammatory tolerance and ageing (Blackwell et al. 1997; Biswas and Lopez-Collazo 2009; Goldberg et al. 2020; Tavenier et al. 2020; Songkiatisak et al. 2022; Nguyen and Cho 2025); our contribution is therefore not the recognition of this phenomenon itself, but the identification of Foxp1-Pde4d and cAMP regulation as a potential mechanism contributing to it. Notably, the Foxp1-dependent recruitment deficit occurred both with and without prior etoposide exposure, indicating that experimentally imposed genotoxic stress was not required for the phenotype. Genotoxic stress therefore provided the discovery context through which Foxp1-Pde4d was identified, whereas tissue injury revealed a broader requirement for Foxp1 in preserving inflammatory responsiveness. The rescue by adenylyl cyclase inhibition provides an important connection between these observations. Reducing cAMP production restored macrophage recruitment in Foxp1-deficient animals, demonstrating that the impaired functional response remained reversible through manipulation of cAMP signalling. Together with the positive regulation of Pde4d by Foxp1, this finding supports a model in which Foxp1-dependent regulation of cAMP contributes to the signalling state from which an inflammatory response is mounted.

A broader role for Foxp1 in injury responses is now emerging across species and tissues. In zebrafish, *foxp1b* mutation increases non-recovery following spinal cord injury (Gillotay et al. 2026). In mice, Foxp1 is upregulated in dorsal root ganglion neurons from three to seven days after sciatic nerve axotomy and is required for sensory axon regeneration, with miR-9 identified as an upstream regulator (Jiang et al. 2017). The convergence of these phenotypes across central and peripheral injury contexts suggests that Foxp1 contributes to successful damage responses in multiple settings.

An important remaining question is where this regulation operates during tissue injury. Foxp1-Pde4d was initially identified in non-immune mesenchymal cells, while *foxp1b*-*pde4d* coupling during zebrafish fin regeneration was concentrated within intermediate epithelial states. One possibility is therefore that Foxp1-dependent cAMP regulation acts within damaged tissue to influence the inflammatory signals that subsequently recruit immune cells, rather than acting exclusively within recruited leukocytes. Such a mechanism would place the regulatory state of tissue-resident cells upstream of immune recruitment, allowing their pre-existing signalling environment to determine how effectively tissue damage is communicated to the immune system. However, the present experiments do not distinguish tissue-resident from immune-cell-autonomous effects, and cell-type-specific manipulation of Foxp1-Pde4d and localisation of the NFκB phenotype will be required to resolve this question.

Altogether, the convergence of single-cell discovery, conserved expression coupling, chromatin regulation, genetic perturbation and in-vivo functional testing identifies Foxp1-Pde4d as a previously unrecognised regulatory axis associated with inflammatory-state heterogeneity and inflammatory responsiveness. Our findings further show that basal inflammatory activity and the capacity to respond to subsequent challenge are related but distinct properties: Foxp1 loss increases basal NFκB activity while diminishing its inducibility. By linking Foxp1-dependent transcriptional regulation to Pde4d and cAMP-sensitive inflammatory function, these findings provide a framework through which cellular state may influence the capacity of inflammatory signalling to adapt as conditions change.

## Methods

### Cell culture and genotoxic stress

3T3-L1 pre-adipocytes were maintained under standard conditions and differentiated using insulin, dexamethasone and IBMX. For genotoxic stress, proliferating cells were exposed to 0.2–0.5 µM etoposide or matched DMSO vehicle for 48 h, followed by drug-free recovery and growth arrest before adipogenic induction. Adipogenic competence was quantified by Oil Red O staining at day 8. Nuclear morphology and Lamin B1, α-tubulin, p53 and Foxp1 abundance were assessed by confocal immunofluorescence, and p53 expression by RT-qPCR. Full culture, staining, imaging and qPCR procedures are provided in Supplemental Methods.

### Bulk RNA sequencing and pathway analysis

Vehicle- and 0.3 µM etoposide-treated cells were collected 4 h after adipogenic induction (n = 3 biological replicates per condition). RNA-seq libraries were sequenced paired-end on an Illumina NextSeq 2000. Reads were processed against the mouse reference genome and differential expression analysed with DESeq2. Pathway and transcription-factor activities were inferred using MSigDB Hallmark gene sets, decoupleR (Badia-I-Mompel et al. 2022), PROGENy (Schubert et al. 2018) and CollecTRI (Müller-Dott et al. 2023). STRING v12 (Szklarczyk et al. 2023) was used to construct functional gene modules.

### Single-cell RNA sequencing and inflammatory-state analysis

Vehicle- and etoposide-treated 3T3-L1 cells were profiled before and 4 h after adipogenic induction using Parse Biosciences Evercode split-pool barcoding. After quality filtering, 3,966 cells were retained and analysed with Seurat v5. Damage-associated trajectories were inferred with Slingshot (Street et al. 2018). IFN/ISG, TNFα/NFκB, senescence and STRING-derived modules were scored per cell. For the transcriptome-wide inflammatory-state screen, expression of each of 7,447 detected genes was correlated with independently defined IFN and NFκB module scores; genes with ρ ≤ −0.10 for IFN and ρ ≥ +0.10 for NFκB defined the IFN-low/NFκB-high quadrant. Functional enrichment used g:Profiler with the screened genes as background.

### Foxp1 ChIP-seq, chromatin interaction and Pde4d analysis

Foxp1 ChIP-seq was performed in duplicate in vehicle- and etoposide-treated 3T3-L1 cells 4 h after adipogenic induction. Peaks were called with MACS2 and reproducible sites defined by IDR; differential occupancy was assessed with DiffBind. Published 3T3-L1 Hi-C and promoter-capture Hi-C data (GSE95533) (Siersbæk et al. 2017) were used to examine contacts from the damage-sensitive Foxp1-bound region at Pde4d, including a distance-matched reflected control. Pde4d transcript abundance and isoform architecture were quantified from GENCODE/Ensembl annotations. Independent Foxp1 gain- and loss-of-function RNA-seq data (GSE179031) (Ma et al. 2023) were re-analysed to test regulation of Pde4d and inflammatory transcriptional programmes.

### Cross-system conservation analysis

Foxp1-Pde4d co-expression was assessed across mouse Tabula Muris Senis (Schaum et al. 2020), human GTEx v8 (GTEx Consortium 2020) and a turquoise killifish multi-tissue atlas (Costa et al. 2026). Conservation was benchmarked against random gene pairs and curated Foxp1 targets. The relationship with inflammatory state was tested in independent single-cell datasets of acute etoposide exposure and replicative senescence (Wechter et al. 2023; Fernandez et al. 2025; Chan et al. 2022). Human adipose single-nucleus data were analysed from Emont et al. (Emont et al. 2022). *foxp1a/b*-*pde4d* coupling during zebrafish fin regeneration was analysed from GSE261907 (Chen et al. 2025), with detection-matched null genes and CS-CORE (Su et al. 2023) used to assess robustness.

### Zebrafish injury and inflammatory responsiveness

Zebrafish were maintained under UK Home Office-approved conditions. F0 *foxp1a/foxp1b* crispants were generated by one-cell-stage CRISPR/Cas9 RNP injection using validated guides (Wafer et al. 2026), with Cas9-only siblings as controls. Tail fins were transected at 3 dpf and larvae imaged longitudinally. Macrophage and neutrophil recruitment were quantified using *Tg(mfap4.1:mTurquoise)* (Walton et al. 2015) and *Tg(lysC:dsRed2)* (Hall et al. 2007) reporters within a 500-µm region proximal to the wound. NFκB activity was measured in a separate *NFκB:eGFP* reporter cohort (Kanther et al. 2011; Feng et al. 2012). To test cAMP dependence, injured larvae were treated with the adenylyl cyclase inhibitor SQ22536; additional pharmacological arms are described in Supplemental Methods.

### Statistical analysis and data availability

Statistical tests were matched to experimental design and are specified in the figure legends and Supplementary Methods. Zebrafish recruitment was analysed primarily by ANCOVA on post-injury values conditioned on the corresponding pre-injury measurement, with experiment included as a fixed block where appropriate. Multiple testing was controlled using Holm or Benjamini-Hochberg procedures as specified. Public datasets re-analysed in this study are listed with their accessions in **Supplementary Table S1**. Analysis code is available at the study GitHub repository (https://github.com/jeminchin/foxp1_etoposide). Sequencing data generated in this study have been deposited at NCBI GEO (accession pending).

## Acknowledgements

We are grateful to Will Cawthorn for comments on the manuscript; Catherine Naughton, Graeme Grimes and Nick Gilbert at the University of Edinburgh for ChIP-seq advice; Yujie Chen, Qinglin Zeng and Ting Wang at Washington University School of Medicine for kindly sharing zebrafish fin regeneration RNA sequencing data; Carl Tucker, Paul Strachan and the BVS aquatics team at the Queen’s Medical Research Institute (University of Edinburgh) for their expert fish husbandry and dedicated care of our zebrafish stocks; and Richard Clark at the Edinburgh Clinical Research Facility for ChIP and RNA sequencing.

## Author Contributions

Conceptualization, JENM, LMHE, PT, YF and RKS; Methodology, LMHE (in vitro / bulk RNA-Seq / ChIP-Seq), PT (scRNA-Seq, zebrafish), GR (zebrafish), LS (zebrafish); Investigation, LMHE (Oil Red O, qPCR, immunofluorescence, bulk RNA-Seq, ChIP-Seq), PT (scRNA-Seq, zebrafish), GR (zebrafish), LS (zebrafish), XC (zebrafish), TM (ChIP-Seq); Formal Analysis, LMHE, PT, JENM; Resources, MNA, YF (zebrafish lines); Writing - Original Draft, LMHE, PT, JENM; Writing - Review & Editing, all authors; Supervision, JENM, PT and RKS; Funding Acquisition, PT and JENM.

## Use of generative AI

JENM used Claude (Anthropic) to assist with drafting text, writing analysis code and analysing existing datasets. All outputs were verified against source data, and the analysis code and rendered notebooks are available at https://github.com/jeminchin/foxp1_etoposide. JENM takes full responsibility for the content.

## Competing Interests

The authors declare no competing interests.

## Funding

The research was funded by a British Heart Foundation PhD Studentship to LMHE (FS/17/62/33477C), a BBSRC research grant to PT and JENM (BB/X009467/1), and a British Heart Foundation REA4 pump-priming award to JENM. YF is funded by a CRUK biology to prevention award (PRCBTP-Nov23/100002).

